# Autophagy controls mucus secretion from intestinal goblet cells by alleviating ER stress

**DOI:** 10.1101/2022.05.26.493604

**Authors:** Maria Naama, Shahar Telpaz, Aya Awad, Shira Ben-Simon, Sarina Harshuk-Shabso, Sonia Modilevsky, Elad Rubin, Jasmin Sawaed, Lilach Zelik, Mor Zigdon, Nofar Fadida, Sondra Turjeman, Michal Werbner, Supapit Wongkuna, Bjoern O Schroeder, Abraham Nyska, Meital Nuriel-Ohayon, Shai Bel

## Abstract

Colonic goblet cells are specialized epithelial cells that secrete mucus to form a barrier between the host and its microbiota, thus preventing bacterial invasion and inflammation. How goblet cells control the amount of mucus they secrete is unclear. We found that constitutive activation of autophagy in mice via Beclin 1 led to production of a thicker and less penetrable mucus layer by reducing endoplasmic reticulum (ER) stress. Accordingly, inhibiting Beclin 1-induced autophagy via Bcl-2 impaired mucus secretion. Furthermore, alleviating intestinal ER stress with a bile acid, or activating the unfolded protein response (UPR) pharmacologically via eIF2α phosphorylation, led to excessive mucus production. Over-production of mucus altered the gut microbiome, with expansion of mucus-utilizing bacteria, and protected from intestinal inflammation. Thus, ER stress is a cell-intrinsic switch that limits mucus secretion, while autophagy maintains proper mucus secretion and intestinal homeostasis by relieving ER stress.

## Introduction

Goblet cells are intestinal epithelial cells that secrete mucus and antimicrobial proteins, creating a chemical barrier between the host and its microbiota. This mucus barrier is crucial for symbiosis between the host and its microbiota as it limits microbial contact with the host epithelium and subsequent proinflammatory responses (Johansson and Hansson, 2016).

Breakdown of this barrier and penetration of bacteria into the inner mucus layer is a hallmark, and perhaps the cause, of inflammatory bowel diseases (IBD) and enteric infections (Johansson et al., 2014; van der Post et al., 2019a). Yet despite their importance, how goblet cells control the amount of mucus they secrete into the gut lumen is unclear.

Macroautophagy (hereafter autophagy) is a cell-intrinsic recycling mechanism crucial for cellular homeostasis which is activated in times of stress such as starvation, infection and accumulation of unfolded proteins in the endoplasmic reticulum (ER). Intestinal secretory cells, such as Paneth and goblet, are highly sensitive to perturbations in the autophagy or the ER stress response pathways. Indeed, mutations in autophagy- and ER stress-related genes pose a risk factor for breakdown of the intestinal barrier and development of IBD. Yet the exact reason for this is not fully clear (Kaser and Blumberg, 2017).

We wanted to understand how goblet cells control the amount of mucus they secrete and why autophagy is needed for proper goblet cell function. Here we show that autophagy relieves ER stress in goblet cells to facilitate mucus secretion. We found that genetic activation of autophagy in mice via the Beclin 1 protein led to suppression of intestinal ER stress which facilitated excess mucus secretion. This over-production of mucus could be achieved, without manipulation of the autophagy process, by pharmacologically reducing ER stress or activating a specific arm of the unfolded protein response (UPR). Excess mucus secretion led to expansion of mucus-utilizing bacteria and protection from colitis. Thus, ER stress is a cell-intrinsic switch that limits mucus secretion, while autophagy maintains proper mucus secretion and intestinal homeostasis by relieving ER stress.

## Results

### Activation of autophagy via Beclin 1 leads to production of a thicker and less penetrable colonic mucus layer

During steady-state the autophagy-initiating protein Beclin 1 is sequestered and inactivated by binding to the B-cell lymphoma 2 protein (Bcl-2). To trigger autophagy, Bcl-2 must be phosphorylated to release Beclin 1 which goes on to initiate synthesis of autophagosomes (Figure 1A) (Fernández et al., 2018). As functional autophagy is vital for proper goblet cell function (Patel et al., 2013), we hypothesized that constitutive activation of the autophagy process will improve goblet cell function and mucus secretion. We thus inspected knock-in mice, in which the phenylalanine residue at position 121 of Beclin 1 is mutated to alanine (*Becn1*^F121A^ mice), preventing binding of Beclin 1 to its inhibitor Bcl-2, leading to constitutive activation of autophagy (Figure 1A) (Fernández et al., 2018). We found that the colonic mucus layer in *Becn1*^F121A^ mice was 50% thicker compared to wild type mice (Figure 1B and (C). To determine whether the thicker mucus layer in *Becn1*^F121A^ mice improved barrier function, we measured the levels of luminal antigens in the sera of mice. Presence of microbial antigens in the blood is directly linked to intestinal barrier function (Serradas et al., 2018). We found lower levels of agonists which activate nucleotide-binding oligomerization domain-containing protein 1 (NOD1), NOD2, toll-like receptors 4 (TLR4) and TLR5 in sera of *Becn1*^F121A^ mice (Figure 1D and E and Figure S1A-C). This implied that the thicker mucus layer of these mice limited translocation of bacterial antigens into host tissues.

**Figure 1.**
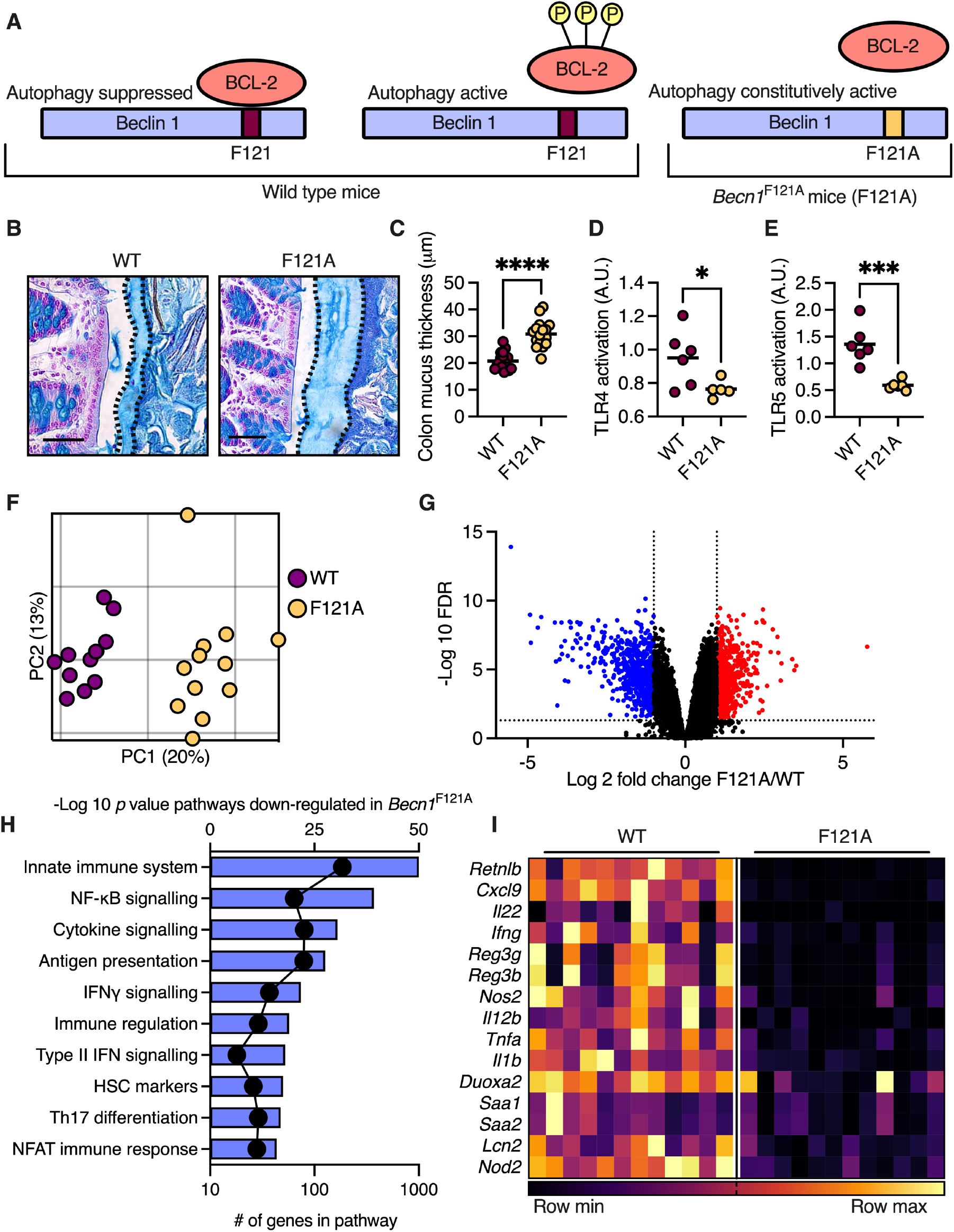
*Becn1*^F121A^ mice produce a thicker and less penetrable colonic mucus layer, and mount a dampened immune response in the colonic tissue. (**A**) Scheme depicting regulation of autophagy activation via Bcl-2 phosphorylation and its alteration in *Becn1*^F121A^ mice. (**B**) Alcian blue staining of Carnoy’s-fixed colonic tissue. The mucus layer is defined by the dashed line. Scale bars, 50μm. (**C**) Measurements of mucus thickness as shown in **B**. (**D**) Detection of TLR4 and (**E**) TLR5 agonist in mouse serum using reporter cell lines. (**F**) Principal coordinates analysis (PCoA) plot of RNA sequencing performed on colonic tissues. (**G**) Volcano plot of transcripts from RNA sequencing. Genes in blue were down-regulated in *Becn1*^F121A^ mice compared with wild type mice and genes in red were up-regulated. (**H**) Pathway analysis of transcripts which are down-regulated in *Becn1*^F121A^ mice according to GO biological function. Bars represent -log (*P* value) and dots represent number of genes in pathway. (**I**) Heatmap depicting differentially expressed innate immune genes with an FDR<0.01. Each column represents a mouse and each row a gene. (**C-F**) Each dot represents a mouse. (**G**) Each dot represents a gene. \**P*<0.05; \*\*\**P*<0.001; \*\*\*\**P*<0.0001; Student’s *t* test. WT, wild type; F121A, *Becn1*^F121A^; A.U., arbitrary unites.

To determine how the improved barrier in *Becn1*^F121A^ mice affected the colonic tissue, we performed a transcriptional analysis. RNA sequencing of colonic tissues from wild type and *Becn1*^F121A^ mice presented a clear difference in transcriptional patterns between the two groups of mice (Figure 1F and G). Unbiased pathway analysis of RNA sequencing data revealed that expression of genes involved in immune regulation and response to bacteria were dampened in *Becn1*^F121A^ mice (Figure 1H). Specifically, levels of transcripts encoding innate immune cytokines, antimicrobial proteins and acute response proteins were diminished in *Becn1*^F121A^ mice (Figure 1I). Thus, constitutive activation of autophagy leads to excess secretion of mucus from colonic goblet cells. This excess of mucus forms a barrier that better limits translocation of luminal antigens into host serum as well as subsequent immune responses in the colon.

### Autophagy reduces ER stress to promote mucus secretion from goblet cells

Next, we wanted to determine the mechanism by which autophagy activation promotes mucus secretion. One possibility is that constitutive activation of autophagy leads to expansion of goblet cell numbers. However, we found a slight reduction in the number of goblet cells in *Becn1*^F121A^ mice compared to wild type mice (Figure 2A). Additionally, transcript levels of transcription factors that mark the goblet cell lineage, *Atoh1* and *Spdef* (Nyström et al., 2021), were similar in the two groups of mice (Figure 2B). Another possibility is that constitutive autophagy promotes mucus secretion by regulating transcription of mucus-forming genes, yet we found no significant differences in transcript levels of these mucus-forming genes when comparing wild type and *Becn1*^F121A^ mice (Figure 2C). Thus, expansion of goblet cells or overexpression of mucus-forming genes could not explain the reason behind the thicker mucus layer in *Becn1*^F121A^ mice.

**Figure 2.**
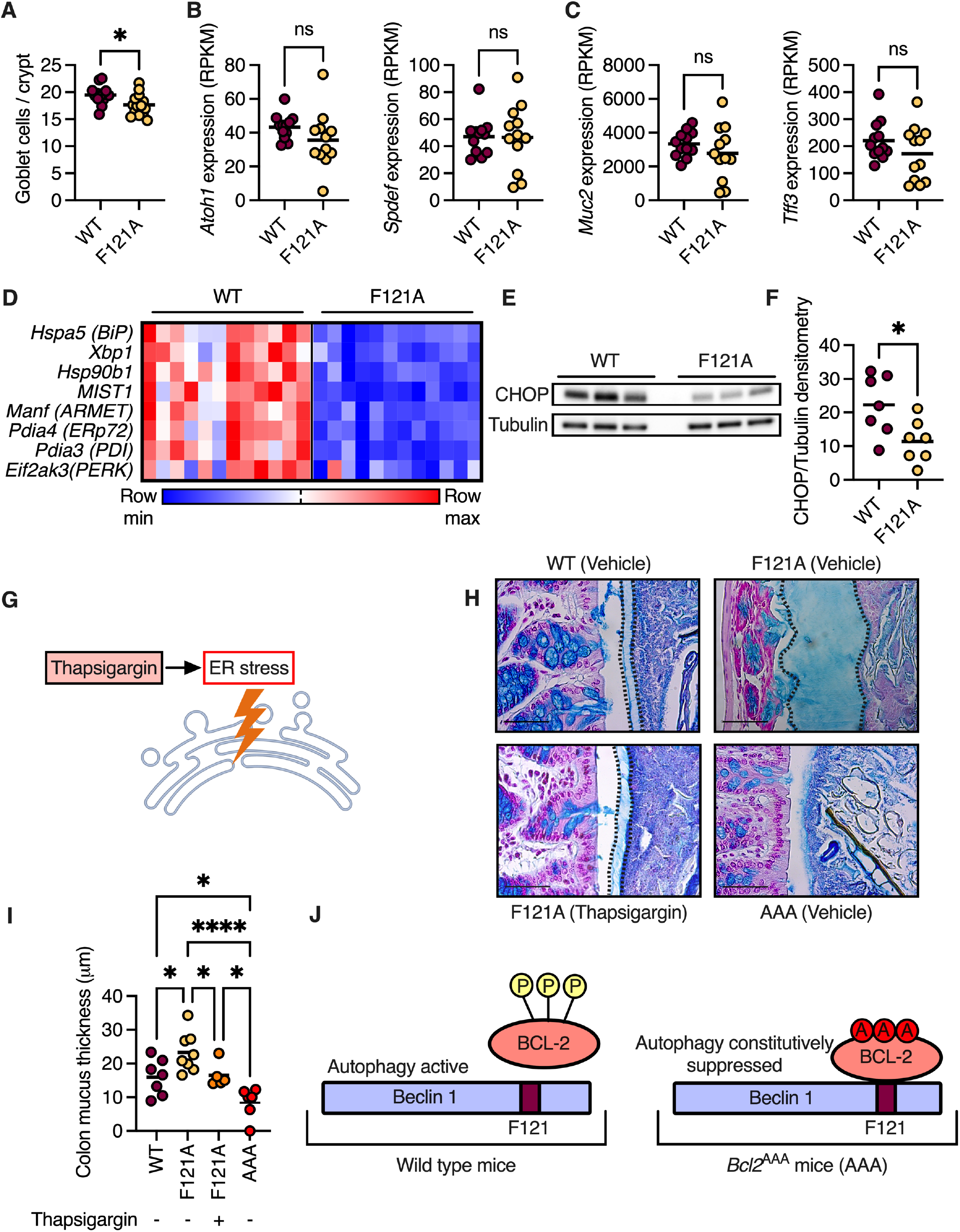
Autophagy relieves ER stress to facilitate mucus secretion from goblet cells. (**A**) Colonic goblet cell numbers per crypt. (**B**) Expression levels of goblet cell-specific transcription factors in colons of mice using RNA sequencing. (**C**) Expression levels of transcripts that encode mucus-forming protein in colons of mice using RNA sequencing. (**D**) Heatmap depicting differentially expressed ER stress response genes with an FDR<0.01. Each column represents a mouse and each row a gene. (**E**) Representative western blot of mouse colons detected with the indicated antibodies. (**F**) Densitometry analysis of western blots as in **E**. (**G**) Scheme depicting ER stress activation via thapsigargin. (**H-I**) Measurements of mucus thickness in colonic section from mice treated with thapsigargin as indicated and stained with Alcian blue. Scale bars, 50μm. (**J**) Scheme depicting regulation of autophagy activation via Bcl-2 phosphorylation and its suppression in *Bcl2*^AAA^ mice. (**A-C, F and I**) Each dot represents a mouse. ns, not significant; \**P*<0.05; \*\*\*\**P*<0.0001. (**A-C, F**) Student’s *t* test; (**I**) One-way ANOVA; WT, wild type; F121A, *Becn1*^F121A^; AAA, *Bcl2*^AAA^. RPKM, Reads per kilobase of transcript.

We then reanalyzed the RNA sequencing data using GO cellular components analysis and found that many of the differentially transcribed genes in the colon of *Becn1*^F121A^ mice encode proteins that are localized to the ER (Figure S2). As a mechanism to relieve ER stress, cells activate the autophagy process (Adolph et al., 2013), while impaired autophagy leads to increased ER stress (Tschurtschenthaler et al., 2017). Of note, impairment of the ER stress response is linked to dysfunction of goblet cells (Kaser and Blumberg, 2009). Indeed, we found that levels of transcripts which are expressed in response to ER stress were reduced in colonic tissue of *Becn1*^F121A^ mice (Figure 2D). Accordingly, protein levels of the ER stress marker C/EBP homologous protein (CHOP) were reduced in colons of *Becn1*^F121A^ mice (Figure 2E and F). Thus, constitutive activation of autophagy reduces ER stress in the colon.

This led us to hypothesize that constitutive activation of autophagy in *Becn1*^F121A^ mice promotes mucus secretion by reducing ER stress in goblet cells. To test this hypothesis, we treated *Becn1*^F121A^ mice with the ER stress inducer thapsigargin (Figure 2G) (Bel et al., 2017). In line with our hypothesis, we found that the mucus layer in thapsigargin-treated *Becn1*^F121A^ mice was decreased compared to that in vehicle-treated *Becn1*^F121A^ mice and was indistinguishable from the mucus layer in wild type mice (Figure 2H and I). To further test the concept that autophagy facilitates mucus secretion by reducing ER stress, we tested whether preventing activation of autophagy via Beclin 1, leaving goblet cells unable to alleviate the ER stress via autophagy, impairs mucus secretion. We used knock-in mice in which the three specific residues of Bcl-2 that are phosphorylated to allow dissociation from Beclin 1 are mutated to alanine (Figure 2J). In these *Bcl2*^AAA^ mice, Beclin 1 is constantly bound to Bcl-2 and thus inactive (He et al., 2012). We found that the mucus layer in *Bcl2*^AAA^ mice was thinner than in wild type mice and in some colonic regions completely absent (Fig. 2H and I). Thus, autophagy facilitates mucus secretion via Beclin 1 by alleviating ER stress in goblet cells.

### Pharmacologically reducing ER stress leads to sustainable excess secretion of mucus

Our observation that activation of autophagy can alleviate ER stress in a manner which results in elevated mucus secretion implies that ER stress might be an intracellular cue that limits mucus secretion. To test this hypothesis, we treated wild type mice with the bile acid tauroursodeoxycholic acid (TUDCA), a chemical chaperone which reduces ER stress (Figure 3A) (Bel et al., 2017). We found that TUDCA treatment resulted in secretion of a thicker mucus layer which resembled the mucus layer in *Becn1*^F121A^ mice (Figure 3B and C). In 20% of TUDCA-treated wild type mice, we found the colonic luminal cavity was filled with mucus (Figure 3D).

**Figure 3.**
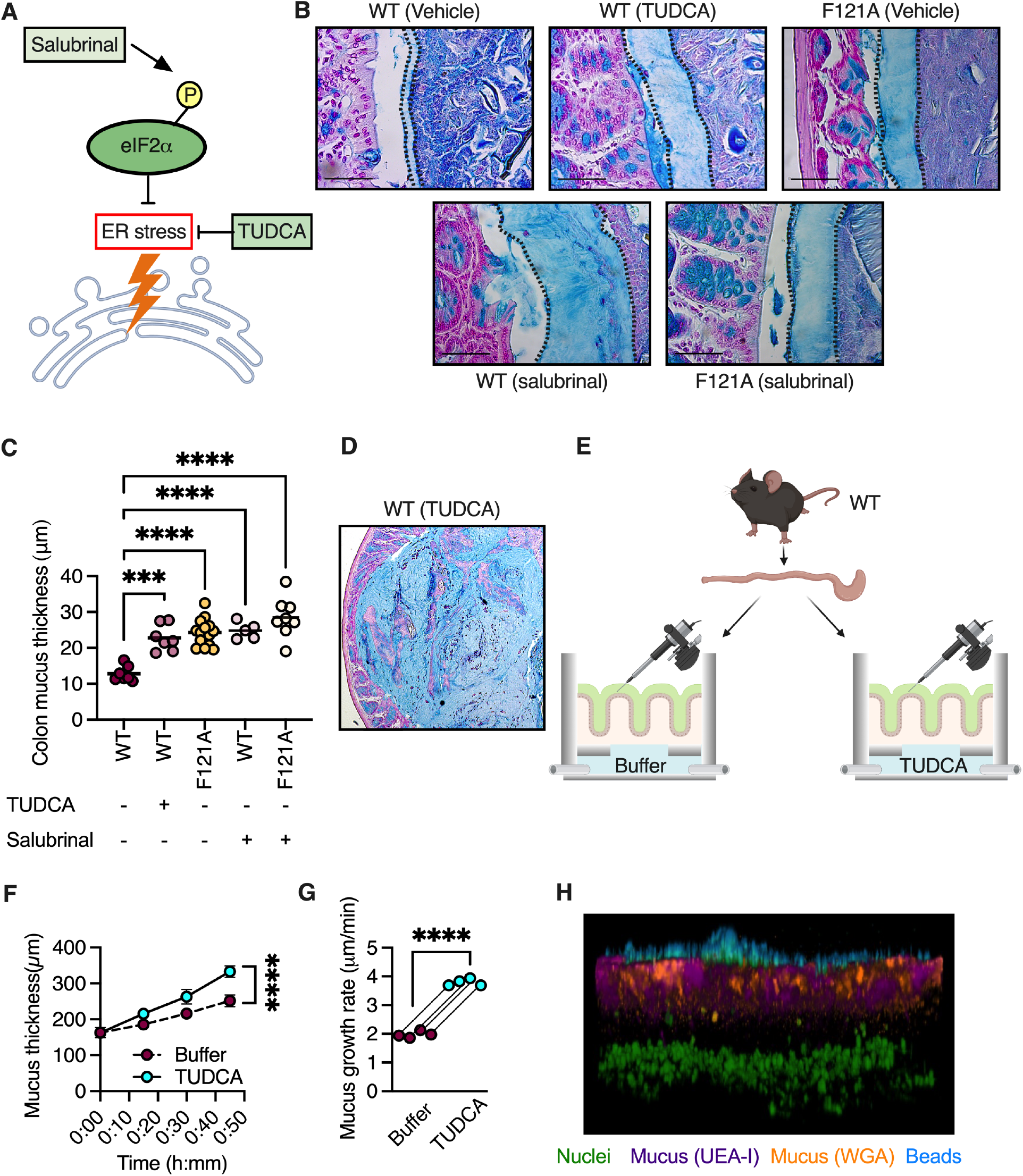
ER stress is a rate limiting switch controlling mucus secretion. (**A**) Scheme depicting ER stress reduction by inhibiting eIF2α dephosphorylation via salubrinal, or by TUDCA. (**B**-**D**) Measurements of mucus thickness in colonic section from mice treated with TUDCA or salubrinal as indicated and stained with Alcian blue. Scale bars, 50μm. Each dot represents a mouse. (**E**) Scheme depicting mucus growth rate measurements. (**F**) Mucus growth over time ±SEM. (**G**) Mucus growth rate. Lines connect tissues from the same mouse. (**H**) Confocal Z-stack imaging of mucus penetrability in TUDCA treated colon explants. Nuclei are in green, UEA-1 lectin in purple, WGA lectin in orange and beads in cyan. \*\*\**P*<0.001; \*\*\*\**P*<0.0001; (**C**) One-way ANOVA. (**F**) Nonlinear regression; (**G**) Paired *t* test. WT, wild type; F121A, *Becn1*^F121A^; TUDCA, tauroursodeoxycholic acid.

TUDCA affects all three arms of the unfolded protein response (UPR) in an unspecific manner (Berger and Haller, 2011). The UPR is a highly regulated cellular program which relieves ER stress to maintain cell function and avoid apoptosis (Kaser and Blumberg, 2009). Thus, we next wanted to determine whether relieving ER stress via activation of a specific arm of the UPR is sufficient for induction of mucus secretion. To test this, we treated wild type and *Becn1*^F121A^ mice with salubrinal, a specific inhibitor of eukaryotic translation initiation factor 2 alpha (eIF2α) dephosphorylation (Figure 3A) (Boyce et al., 2005). The pancreatic ER kinase (PERK) arm of the UPR is activated when the ER stress-sensor PERK phosphorylates eIF2α. When eIF2α is phosphorylated, it inhibits translation of certain transcripts, while promoting translation of transcripts encoding to chaperons and other proteins needed to resolve ER stress (Kaser and Blumberg, 2009). We found that activation of eIF2α in wild type mice via salubrinal treatment led to excessive secretion of mucus from goblet cells, as was seen in TUDCA treated mice (Figure 3B and C). Additionally, the mucus layer in salubrinal-treated *Becn1*^F121A^ mice did not differ from that of vehicle-treated *Becn1*^F121A^ mice (Figure 3B and C), indicating that autophagy activation via Beclin 1 and eIF2α activation converge on the same pathway to induce mucus secretion. Thus, relieving ER stress via activation of eIF2α is sufficient to increase mucus secretion from goblet cells.

We then wanted to determine the kinetics of the elevated mucus secretion that is induced by reduction of ER stress. We removed two adjacent colon sections from each wild type mouse and placed them in two chambers that allow measurement of mucus secretion from live, unfixed tissues under physiological conditions. Only one chamber was infused with TUDCA, allowing us to compare different conditions in tissues that originated from the same mouse (Figure 3E). We found that TUDCA treatment significantly increased mucus secretion rates by two-fold (Figure 3F and G). Additionally, unlike inducing rapid goblet cell degranulation with the cholinergic agonist carbachol, which leads to rapid mucus secretion over 15 minutes followed by reduction of secretion rate (Gustafsson et al., 2012), TUDCA treatment led to a sustained increase in mucus secretion rates over the entire measurement period of 45 minutes (Figure S3). This increase in secretion rates did not impact mucus permeability as compared to the control group (Figure 3H). Thus, ER stress is a cell-intrinsic switch that limits mucus secretion from goblet cells.

### Excess mucus secretion in Becn1^F121A^ mice alters the gut microbiota

The colonic mucus layer also serves as a nutrient source for the microbiota by providing glycans for energy production by bacteria (Hansson, 2020). Thus, we next examined whether constitutive activation of autophagy shapes the gut microbiota via modulation of the mucus layer. 16S rRNA gene sequencing of fecal samples from separately bred and housed mice revealed that *Becn1*^F121A^ mice harbor a distinct gut microbiota compared to wild type mice (Figure 4A and B and Figure S4A), characterized by higher microbial diversity (Figure 4C). Relative abundance analysis revealed that bacteria from the genus *Akkermansia* were enriched in *Becn1*^F121A^ mice (Figure 4D). LEfSe analysis further confirmed that the mucus-utilizing bacterium *Akkermansia muciniphila* was abnormally over-abundant in *Becn1*^F121A^ mice (Figure 4E and F and Figure S4B).

**Figure 4.**
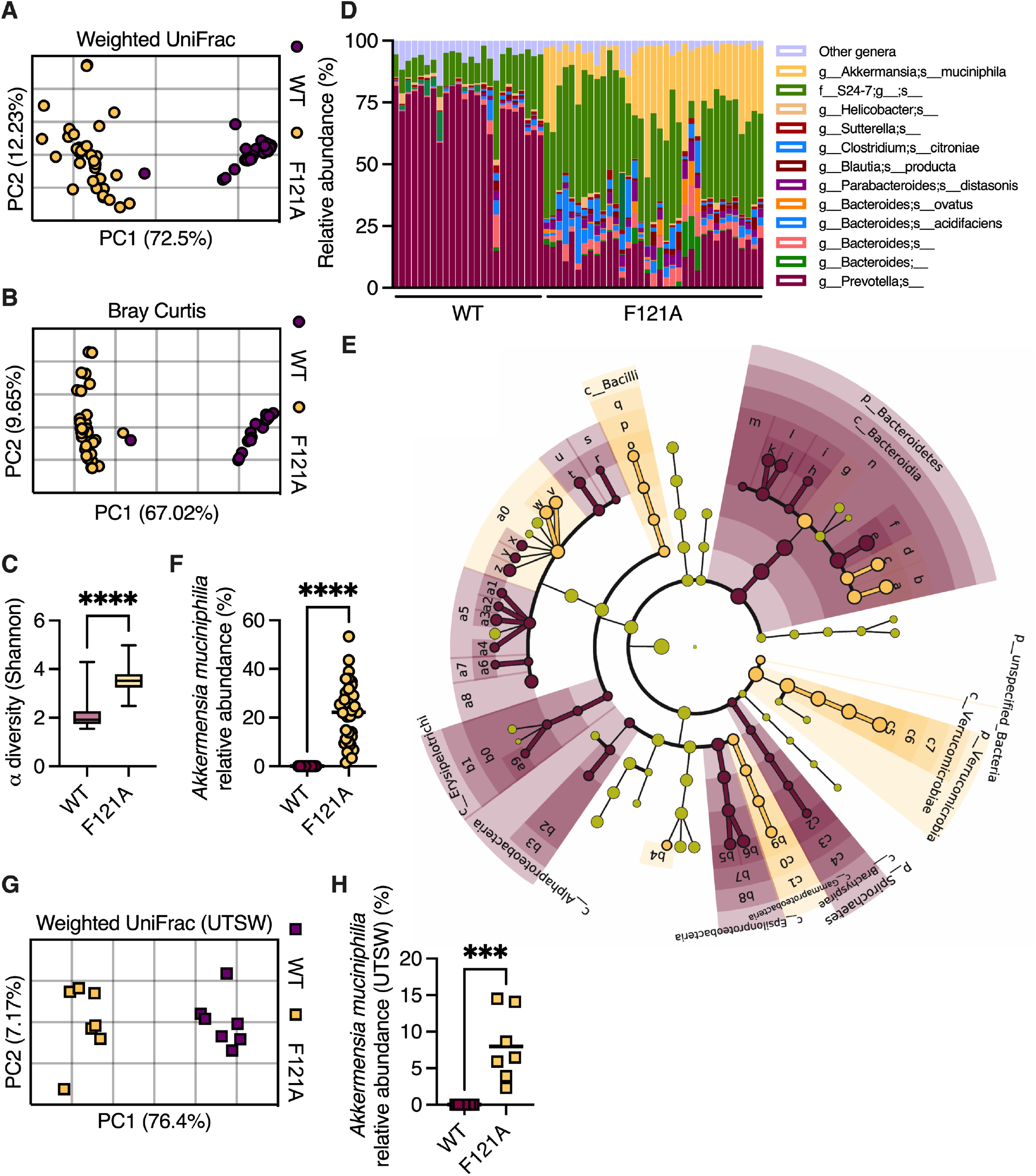
*Becn1*^F121A^ mice contain an altered gut microbiota. 16 S rRNA sequencing was performed to characterize gut microbiota composition. (**A**) PCoA of fecal microbiota β diversity based on weighted UniFrac and (**B**) Bray-Curtis dissimilarity in mice housed at the Bar-Ilan University. (**C**) α diversity comparison based on richness and evenness. (**D**) Relative taxonomic composition. (**E**) Cladogram depicting LEfSe analysis of differently abundant bacteria in wild type and *Becn1*^F121A^ mice. (**F**) Relative abundance of *Akkermansia muciniphila* in mice housed at the Bar-Ilan University. (**G**) Principal coordinates analysis (PCoA) of fecal microbiota β diversity based on weighted UniFrac on mice housed at UTSW. (**H**) Relative abundance of *Akkermansia muciniphila* in mice housed at UTSW. (**A, B, F, G and H**) Each symbol represents a mouse. (**D**) Each column represents a mouse. \*\*\**P*<0.001; \*\*\*\**P*<0.0001; (**C, F and H**) Student’s *t* test. WT, wild type; F121A, *Becn1*^F121A^; UTSW, UT Southwestern.

Variations in microbiota composition are highly dependent on mouse housing conditions and diet (Stappenbeck and Virgin, 2016). To test how conserved the effects of constitutive activation of autophagy are on microbiota composition, we performed 16S rRNA gene sequencing on feces collected from wild type and *Becn1*^F121A^ mice housed at a different animal facility (UT Southwestern Medical Center), fed a diet from a different supplier. Again, we found a clear difference in microbial composition between wild type and *Becn1*^F121A^ mice and an abnormal over-abundance of *Akkermansia muciniphila* (Figure 4G and H). As *Akkermansia muciniphila* is a mucus-utilizing bacterium that can be cultured only on mucus (Desai et al., 2016), its high relative abundance in the colons of *Becn1*^F121A^ mice further confirms the over- abundance of mucus in these mice. Thus, constitutive activation of autophagy reproducibly and rigorously alters the gut microbiota with expansion of mucus-utilizing bacteria.

### The microbiota of Becn1^F121A^ mice confers protects from colitis

Next, we assessed how the altered mucus layer in *Becn1*^F121A^ mice affects susceptibility to colonic inflammation. Penetrability of the colonic mucus layer is directly linked to colonic inflammation in both mice and humans (Johansson et al., 2014; van der Post et al., 2019b). Thus, we hypothesized that *Becn1*^F121A^ mice would be less susceptible to colitis. Dextran sulfate sodium (DSS) treatment instigates colonic inflammation through a direct toxic effect on the mucus barrier (Johansson et al., 2014) and thus was chosen as a suitable model to test the robustness of the mucus layer in *Becn1*^F121A^ mice. When treating mice with 3% DSS for 5 and 7 days we found that *Becn1*^F121A^ mice lost less weight, displayed less signs of severe disease, suffered from reduced colon shortening and had less histological damage as compared to wild type mice (Figure 5A-D and Figure S5A-E). We further confirmed this in a model of moderate colitis by treating mice with 2% DSS for 7 days (Figure S5F-H). Additionally, we found that *Becn1*^F121A^ mice still maintained better barrier function than wild type mice following DSS treatment (Figure S5I).

**Figure 5.**
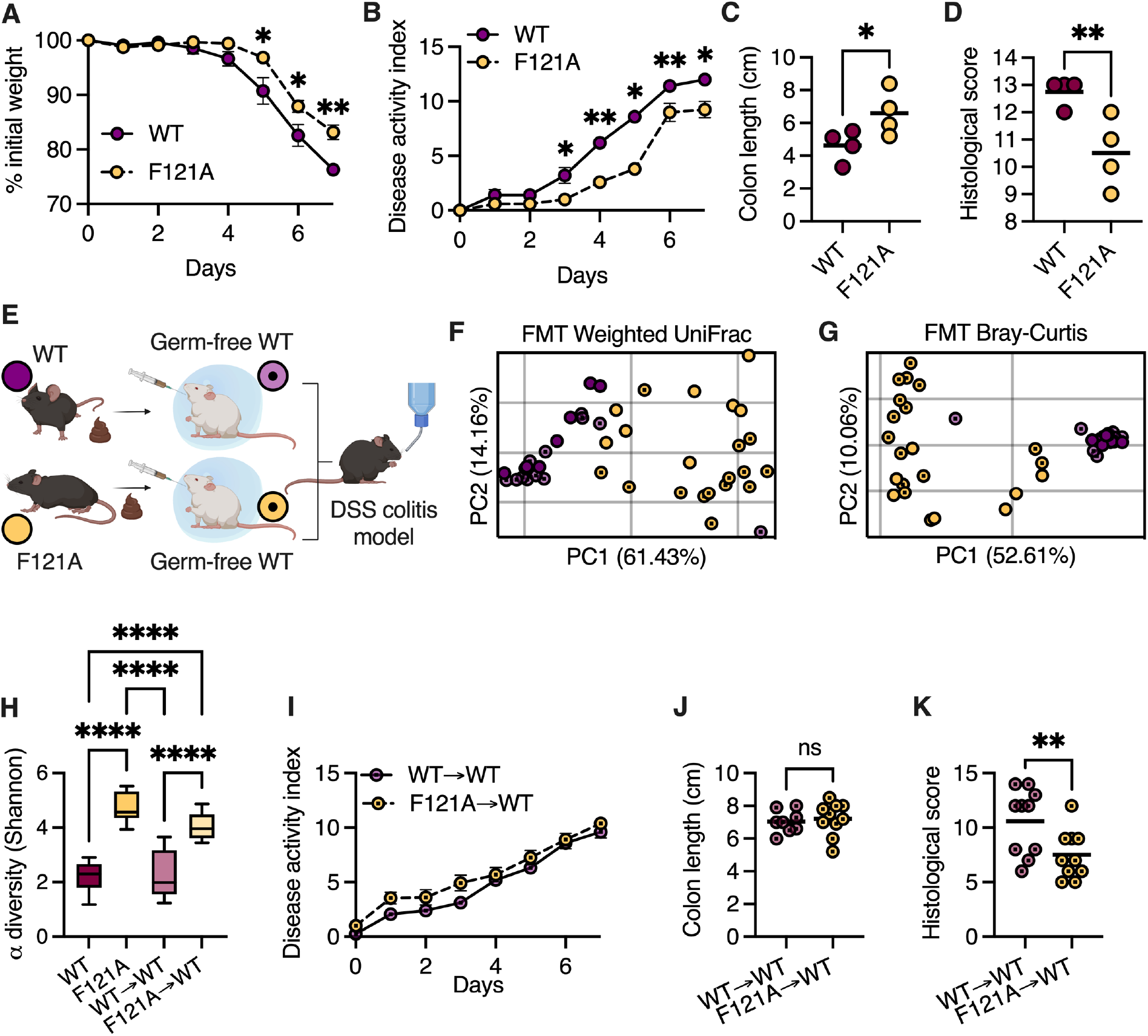
The microbiota of *Becn1*^F121A^ mice confers protects from colitis. (**A-D**) Mice were treated with 3% DSS for 7 days. (**A**) Relative weight change ±SEM, (**B**) disease activity index ±SEM, (**C**) colon length and (**D**) histological damage score. (**E**) Scheme depicting FMT experiment. (**F and G**) PCoA of fecal microbiota β diversity based on weighted UniFrac **(F)** and Bray-Curtis dissimilarity (**G**) of wild type germ-free mice following FMT and separately-housed wild type and *Becn1*^F121A^ mice fecal donors. (**H**) α diversity comparison based on richness and evenness of wild type germ-free mice following FMT. (**I-K**) Mice were treated with 3% DSS for 7 days following FMT. (**I**) Disease activity index ±SEM, (**J**) colon length and histological damage score (**K**). (**C, D, F, G, J and K**) Each symbol represents a mouse. \**P*<0.05; \*\**P*<0.01; \*\*\*\**P*<0.0001; (**C, D, J and K**) Student’s *t* test; (**A, B** and **I**) Multiple *t* tests corrected for false discovery rate; (**H**) One-way ANOVA. WT, wild type; F121A, *Becn1*^F121A^; FMT, fecal microbiota transplant.

The gut microbiota plays an important role in susceptibility to colitis (Bel et al., 2014). As the microbiota of separately bred *Becn1*^F121A^ mice is altered in comparison to wild type mice, we investigated whether this microbiota contributes to the reduced susceptibility of *Becn1*^F121A^ mice to DSS colitis. We colonized germ-free wild type mice with microbiota from either wild type or *Becn1*^F121A^ mice (Figure 5E) and analyzed the effects of fecal microbiota transplant (FMT). While FMT from wild type donors into wild type recipients was highly efficient, FMT from *Becn1*^F121A^ donors into wild type recipients was only partially efficient (Figure 5F and G). This implies that specific conditions present in the intestine of *Becn1*^F121A^ mice are needed to support their microbiota structure. However, the microbiota of recipients from *Becn1*^F121A^ mice was still highly diverse and different than the microbiota of recipients from wild type mice (Figure 5F-H). We then challenged these recipient mice with DSS and found that the microbiota of *Becn1*^F121A^ mice did not prevent weight loss or colon shortening but did reduce histological damage (Figure 5I-K). Thus, the microbiota of *Becn1*^F121A^ mice provides partial protection from colitis.

### Becn1^F121A^ mice are protected from colitis in a microbiota-independent manner

We then investigated whether the distinct microbiota of *Becn1*^F121A^ mice was needed to protect these mice from colitis. We bred heterozygous *Becn1*^F121A^ mice together so that the wild type and *Becn1*^F121A^ offspring were exposed to the same microbiota throughout development and adulthood (Figure 6A). We found that the microbiota of littermate wild type and *Becn1*^F121A^ mice was similar to each other and indistinguishable from the microbiota of wild type mice bred separately (Figure 6B-D). Yet the mucus layer of *Becn1*^F121A^ mice remained thicker than that of their wild type littermates (Figure 6E), indicating that the microbiota did not play a crucial role in promoting excessive mucus secretion in *Becn1*^F121A^ mice. We then challenged these littermate mice, and separately bred wild type and *Becn1*^F121A^ mice, with DSS. We found that *Becn1*^F121A^ mice were still protected from acute colitis, compared to their littermate wild type mice, though to a lesser extent than separately bred *Becn1*^F121A^ mice (Figure 6F and G). Thus, the altered microbiota is not needed for protection against colitis in *Becn1*^F121A^ mice.

**Figure 6.**
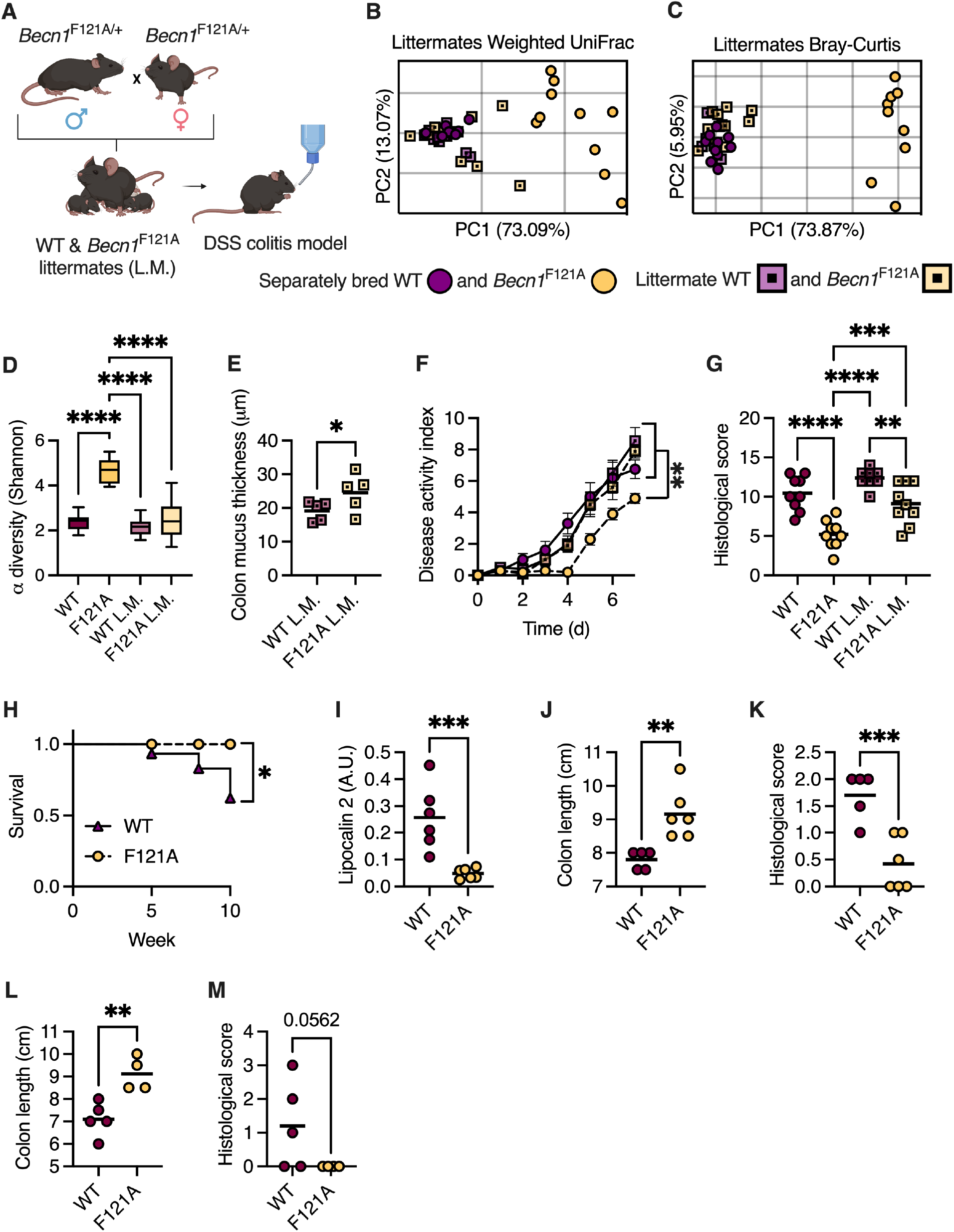
*Becn1*^F121A^ mice are protected from DSS- and AIEC-induced colitis in a microbiota-independent manner. (**A**) Scheme depicting littermate (L.M.) experimental design. (**B**) PCoA of fecal microbiota β diversity based on weighted UniFrac and (**C**) Bray-Curtis dissimilarity of separately bred and littermate wild type and *Becn1*^F121A^ mice. (**D**) α diversity comparison based on richness and evenness. (**E**) Measurements of mucus thickness in colonic section from mice littermate mice stained with Alcian blue. (**F** and **G**) Disease activity index ±SEM (**F**), and histological damage score (**G**) of mice treated with 3% DSS for 7 days. (**H-K**) Survival curve (**H**), stool lipocalin 2 levels (**I**), colon length (**J**) and histological damage score (**K**) of mice pretreated with 20mg streptomycin and infected with 10^8^ C.F.U. AIEC for 10 weeks. (**L** and **M**) Colon length (**L**) and histological score (**M**) of mice pretreated with 20mg streptomycin and infected with 10^8^ C.F.U. AIEC for 14 days. (**B, C, E, G, I-M**) Each symbol represents a mouse. \**P*<0.05; \*\**P*<0.01; \*\*\**P*<0.001; \*\*\*\**P*<0.0001; (**D**, and **G**) One-way ANOVA; (**E, I-M**) Student’s *t* test. WT, wild type; F121A, *Becn1*^F121A^; L.M., littermates; A.U., arbitrary units.

### Becn1^F121A^ mice are protected from infection-driven colitis

Finally, we tested whether *Becn1*^F121A^ mice are also protected from a clinically relevant model of intestinal inflammation by infecting mice with an adherent invasive *E. coli* (AIEC) (Small et al., 2013). AIEC infection is strongly associated with IBD and is thought to be involved in the early stages of IBD development (Buisson et al., 2022). Microbiota differences are negligible in this model as mice are pretreated with antibiotics (Small et al., 2013). We found that, unlike wild type mice, *Becn1*^F121A^ mice did not develop chronic intestinal inflammation or suffer from colon shortening 10 weeks after being infected with AIEC (Figure 6H-K). We further validated these results in an acute infection model (14 days) as well (Figure 6L and M). Thus, constitutive activation of autophagy also protects mice from infection-driven chronic colitis.

## Discussion

Our results illuminate how mucus secretion from goblet cells is controlled by ER stress. As secretory cells tasked with producing large amounts of proteins, goblet cells are inherently susceptible to accumulation of ER stress (Kaser and Blumberg, 2009). Indeed, loss of genes involved in the UPR leads to apoptosis of goblet cells (Kaser et al., 2008). Thus, coupling ER stress to mucus secretion might be a mechanism by which goblet cells preserve viability by reducing mucus secretion when cells are pushed to their limits. Our discovery that autophagy is needed to sustain mucus production by relieving ER stress provides a possible explanation for the prevalence of mutations in key autophagy genes found in IBD patients (Kaser and Blumberg, 2017). With the inability to relieve ER stress because of dysfunctional autophagy, goblet cells might have difficulties keeping up with mucus production demands. This would explain the observation that the mucus layer becomes penetrable in IBD patients (van der Post et al., 2019a) leading to constant contact of microbes with host epithelium and subsequent chronic inflammation.

## Acknowledgments

This study was performed in memory of Dr. Beth Levine, a pioneer and a kind mentor. We thank Lora V Hooper for donating samples from mice housed at UT Southwestern Medical Center.

## Funding

The Azrieli Foundation Early Career Faculty Fellowship (SB)

Israeli Science Foundation (ISF) grants 925/19 and 1851/19 (SB)

European Crohn’s and Colitis Organization (ECCO) Grant (SB)

Mizutani Foundation for Glycoscience (SB)

European Research Council (ERC) Starting Grant 101039927 (SB)

Swedish Research Council (Vetenskapsrådet) – grants 2018-02095 and 2021-06602 (BOS)

## Author contributions

Conceptualization: MN, MW, BOS, MNO and SB

Investigation: MN, ST, AA, SBS, SHS, SM, ER, JS, LZ, MZ, NF, ST, SW, AN and MNO

Funding acquisition: BOS and SB Supervision: SB

Writing – original draft: SB

Writing – review & editing: MW, BOS, ST, MNO and SB

## Competing interests

Authors declare that they have no competing interests.

## Data and materials availability

All data are available in the main text or the supplementary.

## Materials and Methods

### Mice

C57BL/6 wild-type, *Becn1*^F121A^ (Fernández et al., 2018) and *Bcl2*^AAA^ (He et al., 2012) mice were separately bred, unless indicated otherwise, and maintained in the SPF barrier at the Azrieli Faculty of Medicine, Bar-Ilan University, Israel. Germ-free Swiss Webster mice were bred and maintained in isolators as described (Cash et al., 2006). 8–14-week-old mice were used for all experiments. All experiments were performed using protocols approved by the Institutional Animal Care and Use Committees (IACUC) of the Bar-Ilan University.

### Mucus thickness measurements

Mice were euthanized by cervical dislocation, according to IACUC guidelines, and a 1cm long colon tissue sample containing a fecal pellet was removed. Tissues were immediately fixed in Carnoy’s fixative to preserve the mucus layer and processed for paraffin embedding (Johansson and Hansson, 2012). 5μm thick sections were cut using a microtome and stained with Alcian blue stain. Stained sections were visualized using a Zeiss Axioimager M2 microscope and mucus layer thickness (the blue stained area engulfing the fecal pellet) was measured using Zeiss Zen software. 30 measurements were performed per section and the average calculated for each mouse.

### Barrier function analysis

Blood was collected post-mortem via cardiac puncture and incubated at room temperature for 30 minutes in 1.5ml tubes to allow clotting. Samples were then centrifuged at 1,500g for 20 minutes at 4°C after which serum was collected to new tubes. 20μl of serum was added to wells containing InvivoGen HEK-Blue™ reporter cells, and luminal antigens were detected following manufacturer’s instructions.

### RNA sequencing and analysis

RNA from frozen colonic tissues was extracted using Qiagen RNeasy Universal kit. Integrity of the isolated RNA was analysed using the Agilent TS HS RNA Kit and TapeStation 4200 at the Genome Technology Center at the Azrieli Faculty of Medicine, Bar-Ilan University. 1000ng of total RNA was treated with the NEBNext poly (A) mRNA Magnetic Isolation Module (NEB, #E7490L). RNA sequencing libraries were produced by using the NEBNext Ultra II RNA Library Prep Kit for Illumina (NEB #E7770L). Quantification of the library was performed using a dsDNA HS Assay Kit and Qubit (Molecular Probes, Life Technologies) and qualification was done using the Agilent TS D1000 kit and TapeStation 4200. 250nM of each library was pooled together and diluted to 4nM according to the NextSeq manufacturer’s instructions. 1.6pM was loaded onto the Flow Cell with 1% PhiX library control. Libraries were sequenced with the Illumina NextSeq 550 platform with single-end reads of 75 cycles according to the manufacturer’s instructions. Sequencing data was aligned and normalized (reads per million mapped reads) using Partek bioinformatics software. Pathway analysis was performed using the ShinyGO webtool (Ge et al., 2020). Heat maps, principal component analysis (PCA) plots and volcano plots were generated using GraphPad Prism software.

### Immunoblot

Total proteins were extracted from colonic tissue samples by homogenizing in RIPA buffer (Thermo Fisher 89900). 50μg of total protein (as determined by Bradford assay) were loaded onto a 4-20% gradient SDS-PAGE and subsequently transferred to a PVDF membrane. Membranes were blocked with 5% nonfat dry milk in PBS with 0.1% Tween-20. The membranes were incubated at 4°C overnight with the following primary antibodies: α- GADD153/CHOP (SCBT sc-166682) and α-Tubulin (Abcam ab176560). After washing, membranes were incubated with the species-appropriate HRP-conjugated secondary antibodies. Membranes were visualized using an Alliance Q9 system and band density was quantified using FIJI software (Schindelin et al., 2012).

### *In vivo* tauroursodeoxycholic acid (TUDCA), salubrinal and thapsigargin treatment

TUDCA (Sigma T0266), was dissolved in PBS and administered at 250 mg/kg body weight. TUDCA was administered to mice via intraperitoneal injection twice a day for 3 days. On the 4^th^ day, mice were treated in the morning and euthanized 4 hours after the last treatment. Control mice were treated with PBS. Salubrinal (Enzo ALX-270-428) was dissolved in dimethyl sulfoxide (DMSO) at 50 mg/ml and diluted 1:400 in PBS. Mice were treated with 1 mg Salubrinal/kg body weight via intraperitoneal injection once a day for 3 days before being euthanized. Thapsigargin (Sigma T9033), was dissolved in ethanol at 10 mg/ml and then diluted 1:40 in PBS. Thapsigargin (2.5 mg/kg body weight) was administered to mice via intraperitoneal injection once a day for 2 days and euthanized 4 hours after the last treatment. Control mice were treated with ethanol diluted 1:40 in PBS.

### *Ex vivo* mucus measurements and tauroursodeoxycholic acid (TUDCA) treatment

Mucus growth rate in colonic tissue explant was measured as previously described (Gustafsson et al., 2012) with some modifications. Briefly, distal colon tissue from 8-12 week-old wild type mice was collected and washed with 4-5 ml of Kreb’s transport buffer. After muscle removal, tissue was separated into 2 pieces. The tissues were mounted in a heated chamber (37°C). One piece of colon was incubated with RPMI supplemented with 6 mg/ml TUDCA, while another was mounted and incubated with RPMI as a control. Mounted tissues were covered by Kreb’s mannitol buffer, the mucus was overlaid with 10 µm-sized beads and mucus thickness was measured repeatedly with a micromanipulator-connected glass needle. Mucus growth measurement was performed for 45 min.

### Bacterial DNA extraction, amplification, and purification

DNA was extracted from fecal samples using the Invitrogen Purelink™ Microbiome DNA Purification Kit according to the manufacturer’s instructions, following two minutes of bead beating (Bio Spec). Following DNA extraction, the V4 variable region of the bacterial 16S rRNA gene was amplified by polymerase chain reaction (PCR) using the 515F and 806R primers, and each sample received a unique 515F barcoded primer. Primer sequences used were: 515F- (barcode) 5′- AATGATACGGCGACCACCGAGATCTACACGCTAGCCTTCGTCGCTATGGTAATTGTG TGYCAGCMGCCGCGGTAA-3′ and 806 R 5′- CAAGCAGAAGACGGCATACGAGATAGTCAGTCAGCCGGACTACHVGGGTWTCTAAT -3′ (DeSantis et al., 2006).

PCR reactions were carried out with the Primestar taq polymerase (Takara) for 30 cycles of denaturation (95 °C), annealing (55 °C) and extension (72 °C), and a final elongation at 72 °C. Amplicons were purified using AMPure magnetic beads (Beckman Coulter), and subsequently quantified using Picogreen dsDNA quantitation kit (Invitrogen). Samples were pooled at equal concentrations (50 ng/µl), loaded on 2% E-Gel (Thermo Fisher), and purified using NucleoSpin Gel and PCR Clean-up (Macherey-Nagel).

### 16S rRNA gene sequence analysis

Purified products were sequenced using the Illumina MiSeq platform (Genome Technology center at the Azrieli Faculty of Medicine Bar-Ilan University). FASTQ data was processed and analyzed using Quantitative Insights into Microbial Ecology 2 (QIIME2) version 2019.4 (Bolyen et al., 2019). Single-end sequences were demultiplexed, and reads were denoised and clustered using Divisive Amplicon Denoising Algorithm (DADA2) (DeSantis et al., 2006). Primers were trimmed off and single end reads were truncated to ≥ 150 base pairs. A phylogenetic tree was constructed and features were assigned taxonomy using Greengenes reference database with 99% confidence (DeSantis et al., 2006).

Analysis was performed on a rarefied feature table of 7340 reads per sample after filtration of mitochondria and chloroplast sequences. Alpha diversity was calculated using the Shannon index, referring to bacterial evenness within the sample (Pielou, 1966). For between sample diversity (beta diversity), weighted and unweighted UniFrac distances were calculated (Lozupone and Knight, 2005). Over-represented and under-represented features were identified using linear discriminant analysis effect size (LEfSe) (Segata et al., 2011).

### Chemical-induced colitis model

Mice were treated with dextran sulfate sodium (DSS, colitis grade, 36,000 - 50,000 Da, MP) in drinking water at the concentrations and durations indicated in the figure legends. Fresh DSS was prepared daily. Disease activity index (DAI) was measured daily, based on weight loss, stool consistency and rectal bleeding as previously described (Bel et al., 2012). Barrier function using FITC-dextran was performed on the last day as previously described (Bel et al., 2012).

### Infection-induced colitis model

Mice were infected with adherent-invasive *E. Coli* pathotype (AIEC), NRG857c strain (O83:H1). NRG857c was cultured in Luria broth (LB) containing chloramphenicol (34 μg*ml^-^1) and ampicillin (100 μg*ml^-^1). 24h before infection, mice were treated with 20mg streptomycin and then infected with 10^8^ colony forming units (CFU) AIEC. After either 14 days or 10 weeks mice were euthanized, and intestines were fixed in 4% PFA. For quantification of Lipocalin-2 levels, feces were collected and immediately froze in liquid nitrogen and stored at -80°C. Then, samples were reconstituted in PBS containing 0.1% Tween 20, vortexed for 20 min, and centrifuged at 21,000g for 20 min at 4°C. Fecal Lcn-2 levels were determined by the Lcn-2 ELISA kit (R&D systems, DY1857-05) according to the manufacturer’s instructions. Results were calculated by OD and feces weight ratio.

### Histological scoring of colitis severity

Distal colon tissues were fixed in 4% paraformaldehyde, paraffin embedded, sectioned, and stained with hematoxylin and eosin. Histopathological analysis and semi-quantitative scoring were performed by a board-certified toxicological pathologist according to the scoring system described by Cooper *et al*. (Cooper et al., 1993), taking into consideration the grades of extension (laterally, along the mucosa and deep into the mucosa, submucosa, and/or muscular layers) of the inflammation and ulceration. Analysis was performed in a blinded fashion.

### Fecal microbiota transplant (FMT)

Feces collected from mice housed at the SPF barrier facility were immediately transferred into an anaerobic chamber. Feces were vortexed in sterile PBS, debris allowed to settle by gravity, and supernatant transferred to new tubes. Sealed tubes were then removed from the anaerobic chamber and orally administered to germ-free Swiss Webster mice via gavage. Feces from each SPF-housed mouse were used to inoculate two germ-free mice. Inoculated mice were then transferred to Tecniplast IsoCages to prevent outside contamination for two weeks, after which mice were challenged with DSS as above.

## Supplementary Information

**Figure S1.**
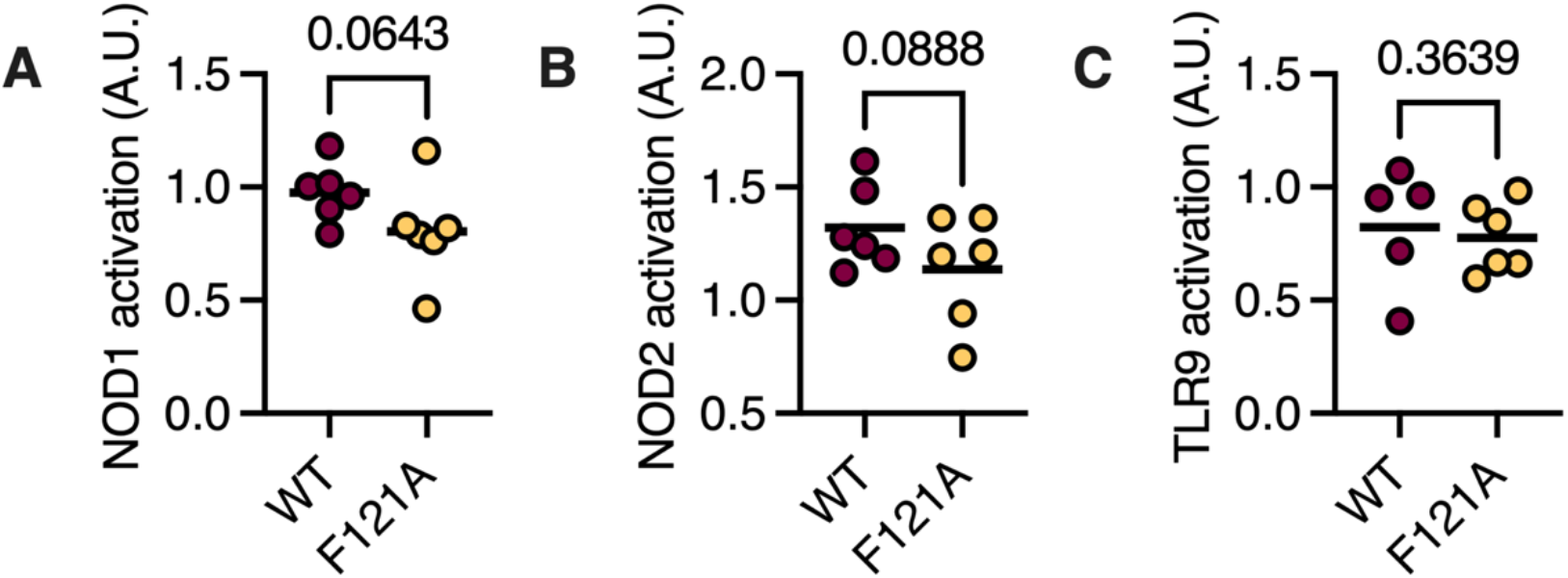
Reduced translocation of luminal antigens into the bloodstream of *Becn1*^F121A^ mice. (A) Detection of NOD1, (B) NOD2 and (C) TLR9 agonist in mouse serum using reporter cell lines. Student’s *t* test; *P* values are displayed. WT, wild type; F121A, *Becn1*^F121A^; A.U., arbitrary unites.

**Figure S2.**
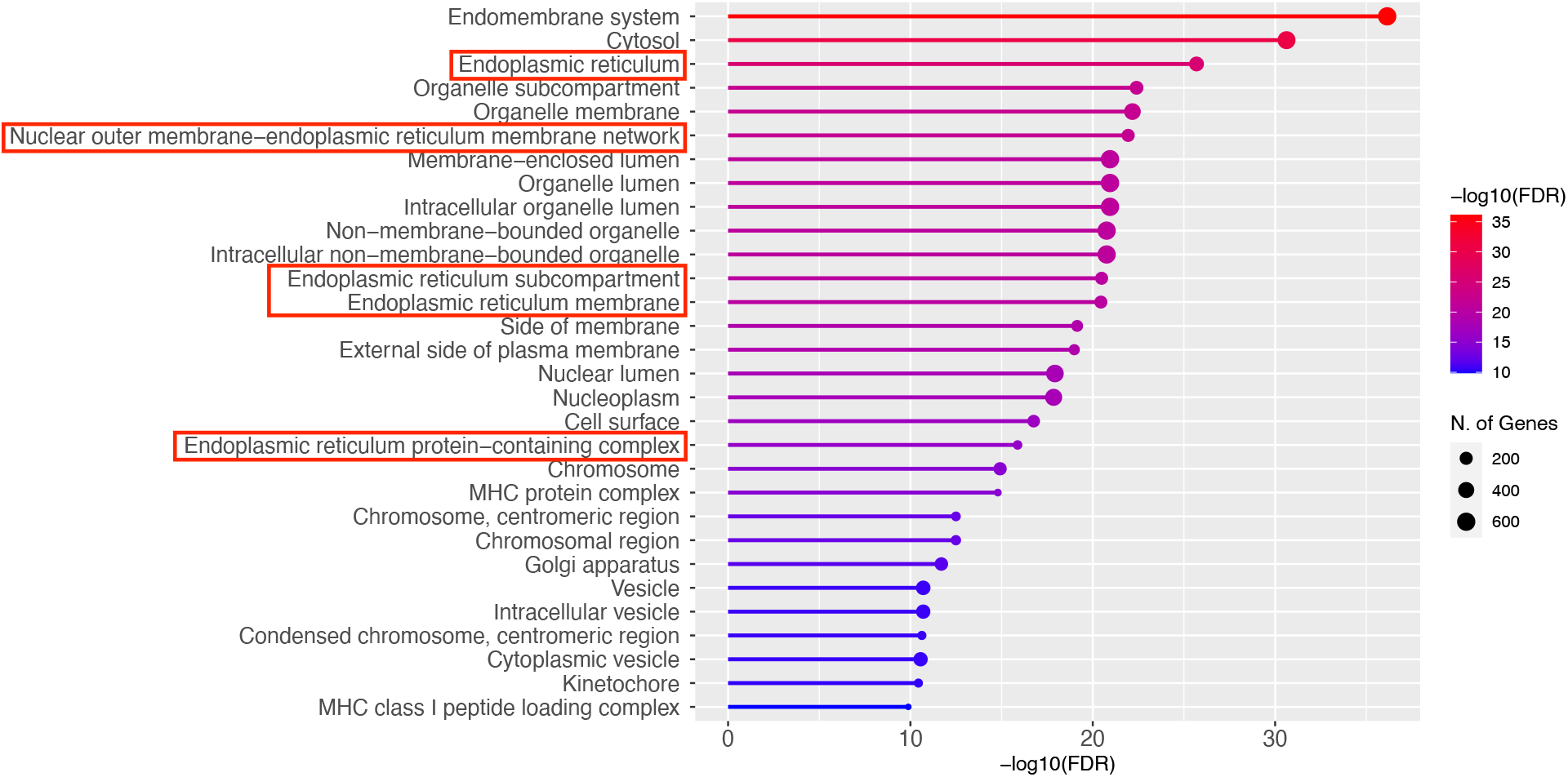
Transcripts of ER proteins are altered in *Becn1*^F121A^ mice. Pathway analysis of differently expressed genes between wild type and *Becn1*^F121A^ mice using GO cellular components analysis. ER-related compartments are marked in red.

**Figure S3.**
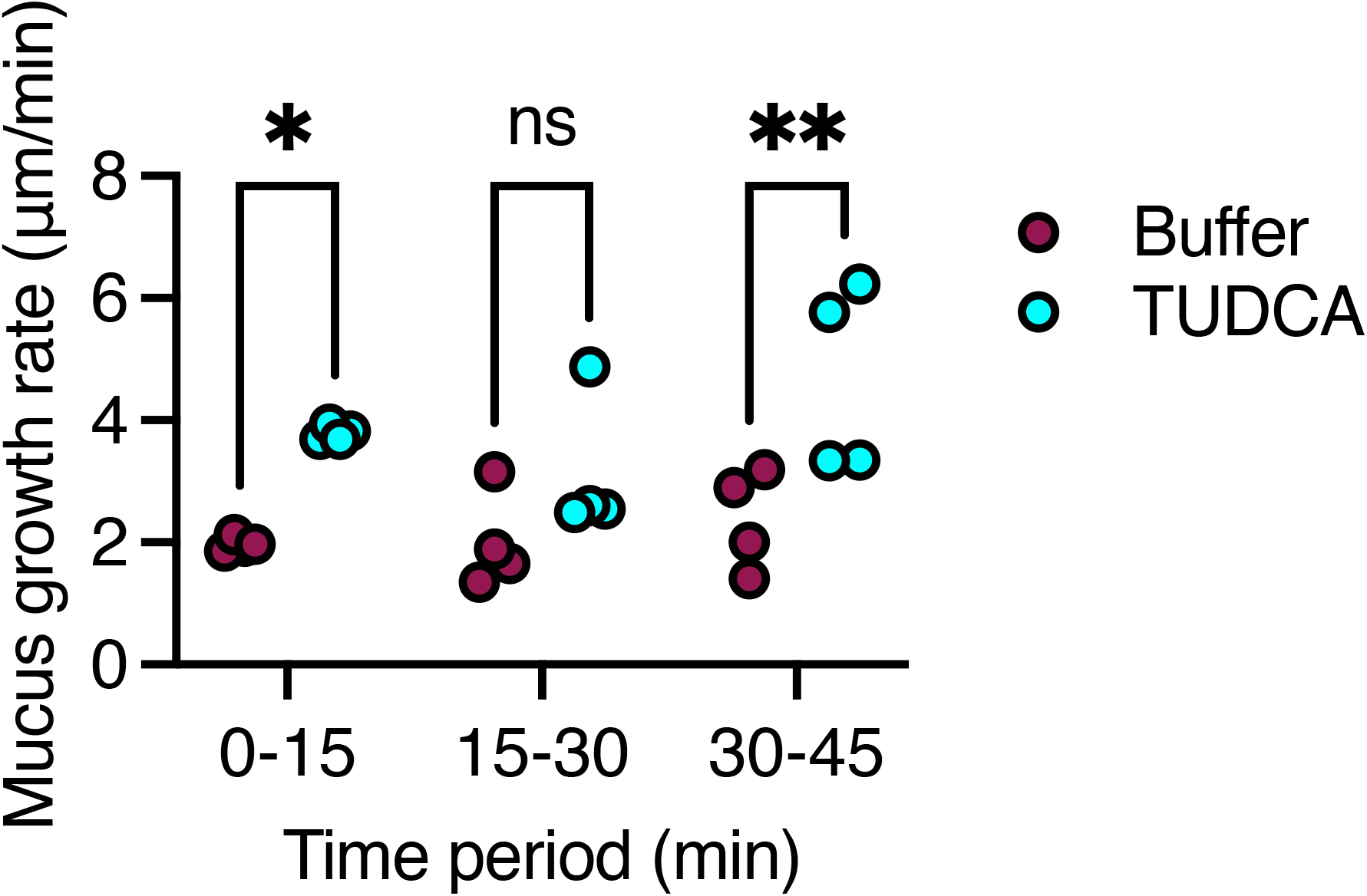
Reducing ER stress increases mucus secretion rate in a sustainable manner. Mucus secretion rates measured in colonic explants treated as noted. Each dot represents a mouse. \**P*<0.05; \*\**P*<0.01; ns, not statistically significant. Two-way ANOVA. TUDCA, tauroursodeoxycholic acid.

**Figure S4.**
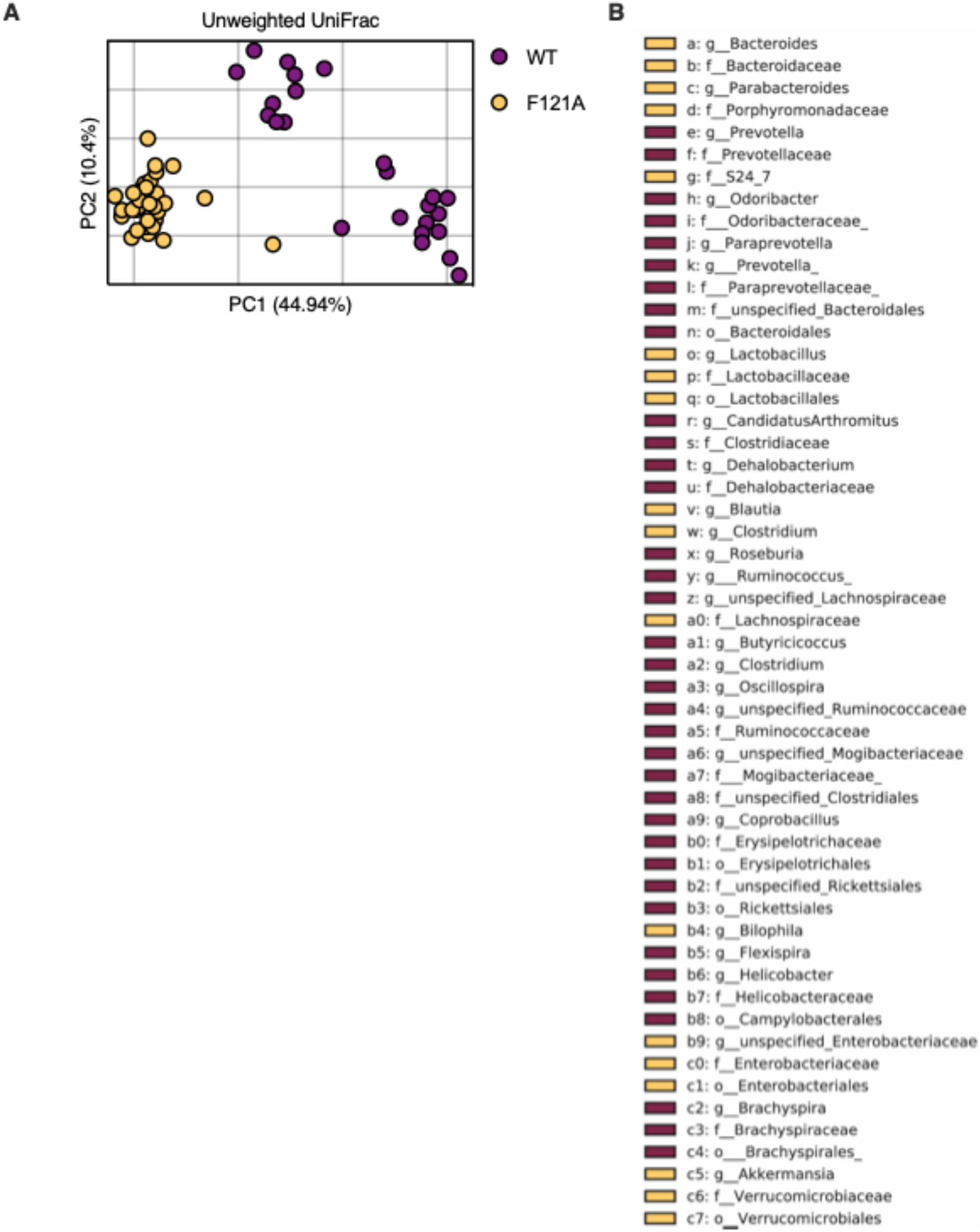
The gut microbiota of *Becn1*^F121A^ mice is different than that of wild type mice and enriched with mucus-utilizing bacteria. 16 S rRNA sequencing was performed to characterize gut microbiota composition. (**A**) PCoA of fecal microbiota β diversity based on unweighted UniFrac. Each dot represents a mouse. (**B**) Legend of cladogram depicting LEfSe analysis of differently abundant bacteria in wild type and *Becn1*^F121A^ mice in **Figure 4E**.

**Figure S5.**
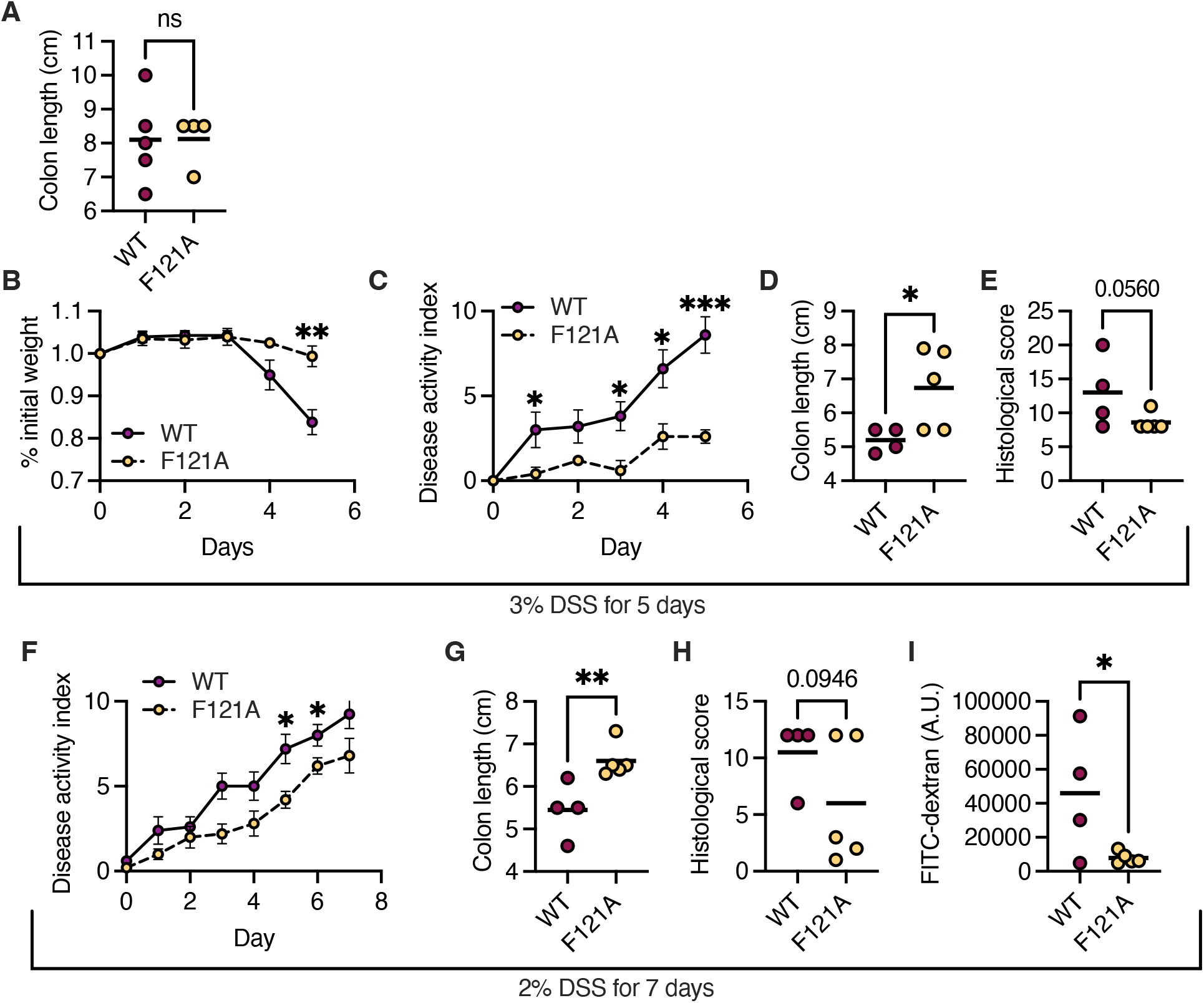
*Becn1*^F121A^ mice are protected from colitis. (**A**) Colon length of naïve wild type and *Becn1*^F121A^ mice. (**B-E**) Relative weight change ±SEM (**B**), disease activity index ±SEM (**C**), colon length (**D**) and histological damage score (**E**) of mice treated with 3% DSS for 5 days. (**F-I**) Disease activity index ±SEM (**F**), colon length (**G**), histological damage score (**H**) and FITC-dextran in serum (**I**) of mice treated with 2% DSS for 7 days. (**A, C-E, and G-I**) Each symbol represents a mouse. \**P*<0.05; \*\**P*<0.01; \*\*\**P*<0.001; (**A, C-E and G-I**) Student’s *t* test; (**B, C** and **F**) Multiple unpaired *t* tests corrected for false discovery rate. WT, wild type; F121A, *Becn1*^F121A^; A.U., arbitrary units.

## Notes

### Competing Interest Statement

The authors have declared no competing interest.

